# *Penicillium purpurogenum* exerts antitumor effects and ameliorates inflammations in Erlich mice model

**DOI:** 10.1101/2020.07.02.184291

**Authors:** Amanda Mara Teles, Leticia Prince Pereira Pontes, Sulayne Janayna Araujo Guimarães, Ana Luiza de Araújo Butarelli, Gabriel Xavier Silva, Flavia Raquel Fernandes do Nascimento, Geusa Felipa de Barros Bezerra, Carla Junqueira Moragas Tellis, Rui Miguel Gil da Costa, Marcos Antonio Custódio Neto da Silva, Fernando Almeida-Souza, Ana Paula Silva de Azevedo-Santos, Maria do Desterro Soares Brandao Nascimento

**Affiliations:** Laboratory for Culture Cell, Universidade Federal do Maranhão, Postgraduate Program in Biotechnology – RENORBIO; Laboratory of Applied Cancer Immunology. Biological and Health Sciences Center, Federal University of Maranhão. Avenida dos Portugueses, 1966, Bacanga. São Luis, Brazil; Immunophisiology Laboratory. Biological and Health Sciences Center, Federal University of Maranhão. Avenida dos Portugueses, 1966, Bacanga. CEP: 65080-805, São Luís, Maranhão, Brazil; Federal University of Maranhão, Postgraduate Program in Adult Health (PPGSAD), Avenida dos Portugueses, 1966, Bacanga. CEP: 65080-805, São Luís, Maranhão, Brazil; Natural Products Department, Institute of Pharmaceutical Techonology, Oswaldo Cruz Foundation, Rio de Janeiro, Rio de Janeiro, Brazil; Centre for the Research and Technology of Agro-Environmental and Biological Sciences (CITAB), University of Trás-os-Montes and Alto Douro (UTAD), 5000-801 Vila Real, Portugal; Post-graduation in Internal Medicine, State University of Campinas, Campinas, São Paulo, Brazil; Post-graduation in Animal Science, State University of Maranhão, São Luís, Maranhão, Brazil; Laboratory of Immunomodulation and Protozoology, Oswaldo Cruz Institute, Oswaldo Cruz Foundation, Rio de Janeiro, Rio de Janeiro, Brazil

**Keywords:** Ehlich’s tumor, *P. purpurogenum*, antitumor, meroterpenoids, inflammation

## Abstract

**Background:** The bioactive metabolites production contributes to the resistance of fungi towards adverse environmental conditions. Some metabolites often have interesting health-promoting activities. This study addressed the anti-tumoural properties of *Penicillium purpurogenum* isolated from a polluted lagoon in Northeastern Brazil.

**Methods:** The extract obtained from the polished environment strain *P. purpurogenum* was fermented, filtered, concentrated and lyophilized, giving rise to the Ethyl Acetate Extracellular Extract (EAE). The metabolites of the extracellular extract of *P. purpurogenum* were studied using direct infusion mass spectrometry. The solid Ehrlic tumor model was used to evaluate the extract antitumor activity. Female Swiss mice were divided in groups (n=10/group) as follow: Negative control (CTL-) treated with phosphate buffered solution; Positive Control (CTL+) treated with cyclophosphamide (25mg/mL); Extracts treatment at doses 4, 20 and 100mg/Kg; Animals without tumor or treatment (Sham); and animals without tumor treated with intermediate dose (EAE20). All treatments were performed intraperitoneally, daily during 15 days. After, the animals were eutanized and the tumor, lymphoid organs and serum were used for immunological, histological and biochemical parameters evaluation.

**Results:** The extract was rich in meroterpenoids. All doses of the extract significantly reduced tumor size compared to CLT- and were associated with 100% survival. Histologically, the 20 and 100mg/kg doses reduced tumour-associated inflammation and tumour necrosis. The extract also reduced cellular infiltration of lymphoid organs and circulating TNF-α levels when compared with CLT-. The extract did not induce weight loss and renal or hepatic toxic changes.

**Conclusions:** These results indicate that *P. purpurogenum* from a polluted marine environment produce hybrid natural products of the terpenoid pathway that exhibits immunomodulatory and antitumor properties *in vivo*. Thus, fungal fermentation is a biotechnological approach for the production of antitumour agents.

## BACKGROUND

Fungi are versatile organisms that can be found in several habitats, occupying inhospitable ecological niches in all ecosystems on the planet, with promising therapeutic and biotechnological potential [1]. Several studies show that marine microorganisms are sources of unique natural products including molecules with potential anticancer uses [2,3,4,5].

*Penicillium* fungi synthesize high amounts of known bioactive secondary metabolites [6,7] including anticarcinogenic drugs and immunosuppressive agents [8,9]. *P. purpurogenum* species has the ability to synthesize a variety of substances with biotechnological and bioactive potential, but the strains that demonstrate this activity are mutant strains resistant to antibiotics [10,11].

*P. purpurogenum* MA52 is a strain previously isolated from a polluted marine environment, the Jansen lagoon in the Brazilian Northeastern state of Maranhão [12]. This is a highly pollutes environment receiving domestic effluents from the surrounding city of São Luís [12]. However, the specific characterisitcs of the MA52 strain which allow it to survive in this polluted environment have not yet been described. New drugs are being developed from secondary metabolites in search of less toxic and more effective compounds compared with traditional cancer therapies [13]. Also, some natural products become important nutraceutics with cancer chemopreventive properties [14,15].

Many *in vivo* models are used for studying breast cancer, the most common cancer in women worldwide [16]. Breast cancer models include genetically-modified animals, xenografted tumours in immune-compromised mice, chemically induced models [17], and syngeneic models like the solid Ehrlich’s tumor, a spontaneous and highly aggressive murine mammary adenocarcinoma [18]. This syngeneic model is immunocompetent and avoids the use of harmful chemicals, making it particularly useful for tumor and chemotherapy studies [19,20]. Considering the search for bioactive compounds capable of being prototypes of new antitumor drugs, the present study aims to describe the *in vivo* antitumor activity of *P. purpurogenum* extract compounds obtained from the MA52 strain against solid Ehrlich’s tumor.

## METHODS

### Fermentation and Preparation of Ethyl Acetate Extracellular Extract (EAE)

*P. purpurogenum* is a marine fungus found in the coastal region (2°29’56”S 44°17’59”W) of Maranhão, Brazil. The present strain was isolated by the Mycology Laboratory of the Basic and Applied Immunology Center of the Federal University of Maranhão and deposited in the fungi collection of the Federal University of Maranhão under access code MA52. Fungus strain was grown in Potato Dextrose Agar (BDA) at 28 °C for 7 days until complete growth. After that period, superficial circles of mycelium-containing agar were further cultivated in BDA broth for fermentation at 28 °C for 21 days in a rotary shaker at 150 rpm. Afterwards, 300 mL of the fermented broth were macerated for 48 hours with 600 mL of ethanol, to obtain an extracellular extract. Then, aqueous ethanol solution was filtered and concentrated under reduced pressure to remove ethanol and the remaining water was extracted twice with 1:4 (v/v) ethyl acetate, resulting in an organic phase which was concentrated and lyophilized to obtain the ethyl acetate extract used for *in vivo* testing.

### Tandem Mass Spectrometry with Electrospray Ionization (ESI-MS/MS)

Ethyl acetate extracellular extract (EAE) was analyzed by direct infusion (ESI-MS/MS) in a Bruker Ion trap amazon SL mass spectrometer, positive mode (ESI+). EAEE (3mg) was dissolved in methanol certified HPLC grade containing 0.1% formic acid (v/v) using an ultrasonic bath for 20 minutes. The operating conditions were 1 □L/min infusion, 4.0 kV capillary voltage, 100 °C temperature source, and cone voltage of 20-40 V. Mass spectra were recorded and interpreted by Bruker Compass Data Analysis 4.2.

### Animals

After approval by the Ethics Committee (CEUA 23115.11239/2017-70), female Swiss mice (n=70), weighing 25-30 g and aged between 3 and 4 months, were provided by the Federal University of Maranhão (UFMA). Animals were kept in room with controlled temperature at 22 ± 3 °C, 50 ± 15% relative humidity, 12 hours light/dark photoperiod, and food and water *ad libitum*. Animal experiments were conducted according to animal welfare guidelines by the National Council for Control of Animal Experimentation (CONCEA). Every effort has been made to reduce the number of animals used and their discomfort.

### Ehrlich solid tumor model

Animals were anesthetized with ketamine/xylazine (120-150 mg/kg). Ehrlich ascitic carcinoma was maintained in mouse via intraperitoneal injections of 2×10^6^cells [21]. For inducing the solid tumor, transplantable neoplastic cells with 7 days of ascitic evolution were aspirated and 200 µL of cell suspension at 2×10^6^ cells/mL was injected into the left posterior foot pad. The cells were found to be more than 99% viable by a Trypan blue exclusion method. The experimental treatment was started 24 hours after the tumor implantation [22].

### Treatment Groups

The animals were separated in two groups: with tumor inoculation for antitumor activity and without tumor inoculation used to evaluated the extract toxicity. The tumor inoculation group was subdivided in six subgroups (n=10), the negative control subgroup (CTL-) with tumor induction treated with phosphate buffered solution (PBS); the positive control subgroup (CTL+) treated with cyclophosphamide at dose of 25 mg/kg; and the subgroups treated with extracts at doses of 4 mg/kg (EAE4+Tumor), 20 mg/kg (EAE20+Tumor) and 100 mg/kg (EAE100+Tumor). To determine acute toxicity, two groups without tumor induction were treated with PBS (Sham) or with the extract at medium dose of 20mg/kg (EAE20). All treatments were performed intraperitoneally 24 hours after tumor inoculation during fifteen days, the volume admistrated was 100µL. After treatment, the animals were randomly selected, a part was euthanized (n=5) for biological analysis and another part (n=5) was used for the quality of life/survival test. The euthanasia was performed by the administration of 120 mg/kg ketamine and 150 mg/kg xylazine (2:1 solution) via intraperitoneal [23] and underwent complete necropsy.

### Tumor development Assay

The Ehrlich solid model was performed to evaluated antitumoral activity. Paw volume was determined before and after the injection of Ehrlich tumor cells using a digital caliper at 48 h intervals. The volume was calculated by multiplying the thickness, width and length measurements of the paw with tumor presented as mm^3^. After the treatment performed in fifteen days, the animals of each group were randomly chosen (n=5), euthanised and the paws with a tumor were removed and weighed. The area under the curve was calculated from the kinetics graph obtained from the tumor-inoculated paw development data using the GrafPAd Prism 7 software.

### Histopathological analyzes

Immediately after euthanasia, Ehrlich’s tumor tissue (and matched normal tissue from sham groups), liver and kidneys were removed and fixed in 10% neutral buffered formalin. Samples sections were stained with hematoxylin and eosin solution and analyzed by a single researcher with expertise, blinded to the experimental groups. Concerning the tumour samples, the following parameters were analyzed: inflammatory infiltrate distribution (focal, multifocal, diffuse or peripheral); inflammatory infiltrate intensity (scores: absent 0, light 1, moderate 2, intense 3); necrosis (scores: absent 0, focal 1, focally extensive or multifocal 2, diffuse 3); mitotic figures (scores: no mitotic figures 0, occasional mitoses 1, single mitotic figure per 400x field 2, two mitotic figures per 400x field 3); cellular pleomorphism (scores: monomorphic tumour cells 0, minimal intercellular variation 1, variations in nuclear size and shape 2, major variations with bizarre nuclei 3); and tumour invasion (scores: well-defined borders with no obvious invasion 0, well-defined borders with minimal invasion ofadjacent tissues 1, poorly-defined borders with marker invasion 2, unrecognizable borders with multiple tumour foci 3) [24].

### Lymphoid organs cellularity

To obtain and quantify cells in the popliteal lymph node and spleen, these solid organs were removed and macerated in 1mL PBS. To obtain bone marrow cells, the femur was removed and perfused with 1 mL of PBS. After, 90 μL of lymph node, spleen and bone marrow cell suspensions were added to 10 μL of violet crystal and the cells were counted in a Neubauer chamber with the aid of a common light optical microscope [25].

### Blood samples

Blood samples were obtained by cardiac puncture. The samples were then centrifuged at 5000 rpm for 10 min. Serum were separated and stored in aliquots at - 80°C until needed. Prior to the assay, the samples were thawed at room temperature [23, 24].

### Cytokine Quantification

Blood serum was used for quantification of interleukin-2 (IL-2), interleukin-4 (IL-4), interleukin-6 (IL-6), interferon-γ (IFN-γ), tumor necrosis factor (TNF-α), interleukin 17A (IL-17A) and interleukin-10 (IL-10) by flow cytometry with FACSCalibur equipment (BD Biosciences, San Jose, CA, USA), using the BD ™ Cytometric Bead Array (CBA) cytokine kit Mouse Th1/Th2/Th17 (BD Biosciences, San Jose, CA, USA) following the manufacturer’s recommendations.

### Extract toxicity

For the assessment of acute toxicity, we take into account the weight of the animals during the treatment, weighed daily for 15 days. Weight variation was verified before the tumor inoculation until the last day of treatment. We also verified that it is a fungus extract, hepatic parameters in which blood serum was used to perform the biochemical measurement of glutamic-oxalic transaminase (TGO), glutamic-pyruvic transaminase (TGP) through colorimetric analyzes using the Labtest kits, following the manufacturer’s guidelines, and the data was obtained on a visible spectrum plate reader (Lab. Syftemf Multi Skan EX). Histopathological analysis of the liver and kidney was performed. The parameters analyzed in the liver were the presence of mitotic figures, caryatia (more than 10% of hepatocytes with nuclei twice the size of normal hepatocytes) and necrosis present or absent. Hepatitis was classified as mild (hyperplasia of Küpfer cells and/or occasional microabscesses or mild focal periportal leukocytic infiltration) or moderate (multifocal to diffuse leukocytic infiltration in multiple portal spaces or centrilobular veins). Hepatocellular vacuolar degeneration was also classified as mild (restricted to the periportal and/or centrilobular areas) or moderate (extending to the midzonal areas). In the kidney, the presence of tubular degeneration, defined as the swelling of the cells of the outlined proximal or distal tubules, or necrosis of isolated tubular cells or loss of cell vesicles in the tubular lumen (bleeding) was evaluated [26]. The quality of life/survival of the animals was verified for 30 days after treatment, following a quality of survival protocol in which they checked some of the animals that suffered damage, and a pain scale, in which if the animal shows three or more potential signs associated with pain or discomfort they are kept under surveillance and staying for more than 72 hours, being euthanized by excess of anesthetic [27].

### Statistical analysis

Results were expressed as mean ± standard error of means (S.E.M. and S.D.). The differences were submitted to analysis of variance (one-way or two-way ANOVA) followed by Newman-Keuls test and by Student’s t-test using GraphPad Prism software, version 7.0. To evaluate the survival curve, the Kaplan-Meier curve was used and the statistical analysis was performed by the Log-Rank test. The significance level for rejection of the null hypothesis was 5% (p <0.05).

## RESULTS

### *P. purpurogenum* extracellular extract contains meroterpenoids

The mass spectral data obtained by direct infusion in positive mode (ESI-MS/MS), presented at figure 1, were useful to determine the chemical profile of *P. purpurogenum* extracellular extract (EAE) and showed the presence of five major compounds (1-5). They were tentatively identified as meroterpenoids, compounds commonly produced by fungi [28], by comparing the mass spectral data with that described in the literature.

**Fig 1.**
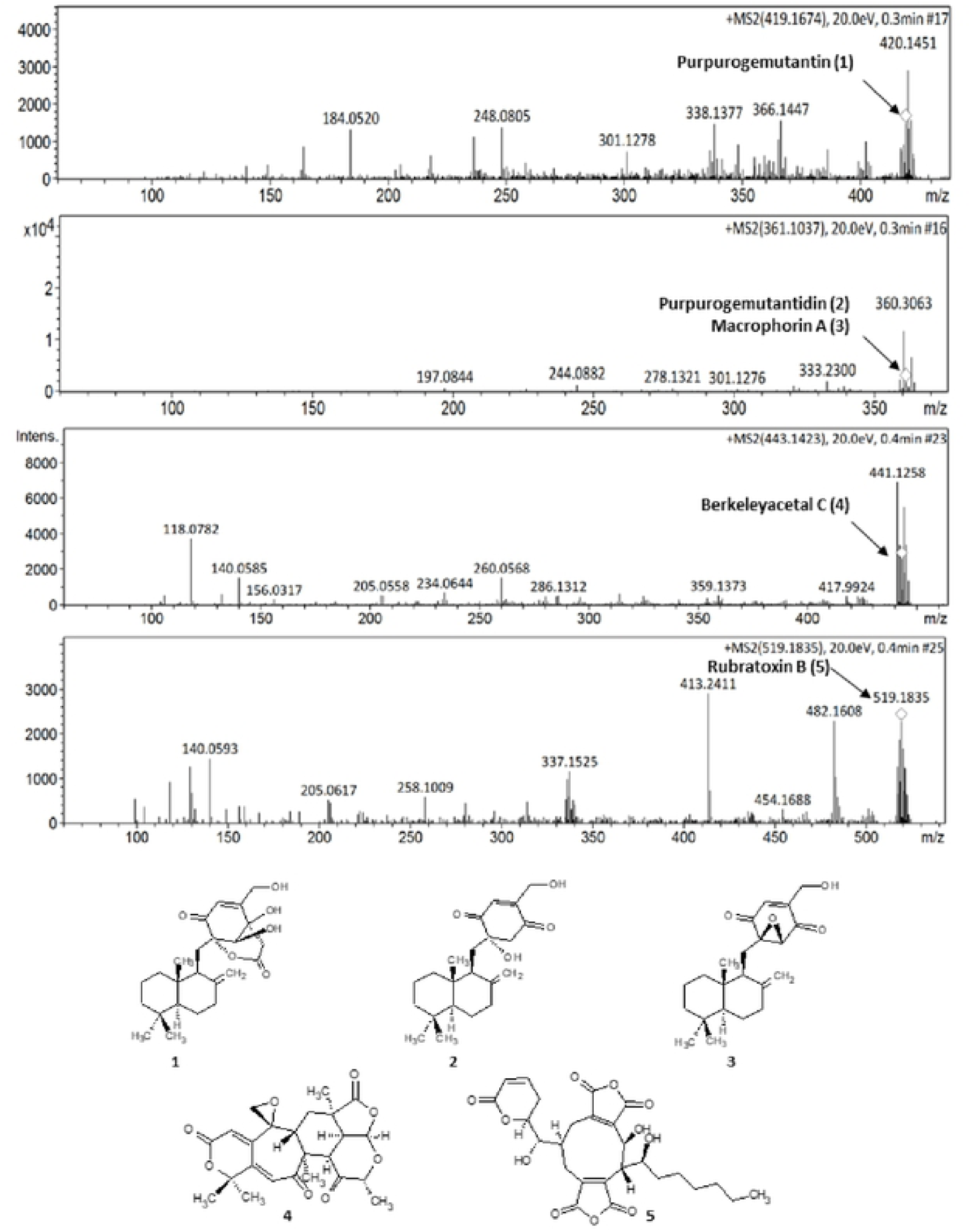
Mass spectra obtained for tentative identification of meroterpenoids 1-5.

Compound **1 (**purpurogemutantin), showed experimental pseudo-molecular ion (M + [H]^+^) at *m/z* 419,1674 compatible with the molecular formula C_24_H_35_O_6_. Compounds **2** (purpurogemutantin) and/or **3** (macrophorin A) showed the same pseudo-molecular ion (M + [H]^+^) at *m/z* 361.1037 corresponding to the molecular formula C_22_H_33_O_4._ These compounds were previously described at *P. purpurogenum* extracts by Fang et al [29]. **4** (Berkeleyacetal C, molecular formula C_24_H_26_O_8_), another meroterpenoid previously described in mutant *P. purpurogenum* by Li et al [30] presented pseudo-molecular ion (M + [H]^+^) at m/z 443,1696 while compound **5** (Rubratoxin B) showed pseudo-molecular ion (M + [H]^+^) at *m/z* 519.1758 corresponding to the molecular formula C_22_H_33_O_4_. Rubratoxin B is a known meroterpenoid with anti-cancer activity isolated by *P. purpurogenum* [31].

### *P. purpurogenum* extract showed activity against Ehrlich’s solid tumor

In tumor growth kinetics, the results showed that throughout the experiment the negative control group presented an upward curve, while those treated with cyclophosphamide and the extract showed a linearity kinetics. The extract significantly reduced tumor growth from the eighth day compared to CLT-, while cyclophosphamide reduced the tumor growth from the tenth day (Figure 2a). The kinetic values of tumor growth have been in accordance with the graph of the area under curve (AUC). The extract at doses 4 mg/kg (422±35.8mm^2^), 20mg/kg (389.7±49.13 mm^2^) and 100mg/kg (365.5±40.47mm^2^) presented an area smaller than the negative control (1240±43.25 mm^2^) and the chemotherapy (599.8±55.51mm^2^) (Figure 2b). At the end of the experimental period, the extract at 4mg/kg (0.19±0.02g), 20mg/kg (0.19±0.05g) and 100mg/kg (0.16±0.02g) showed low weight of the tumor when compared to the negative control (0.38±0.07g) and showed similar weight to the positive control (0.15±0.05g) (Figure 2c).

**Fig. 2.**
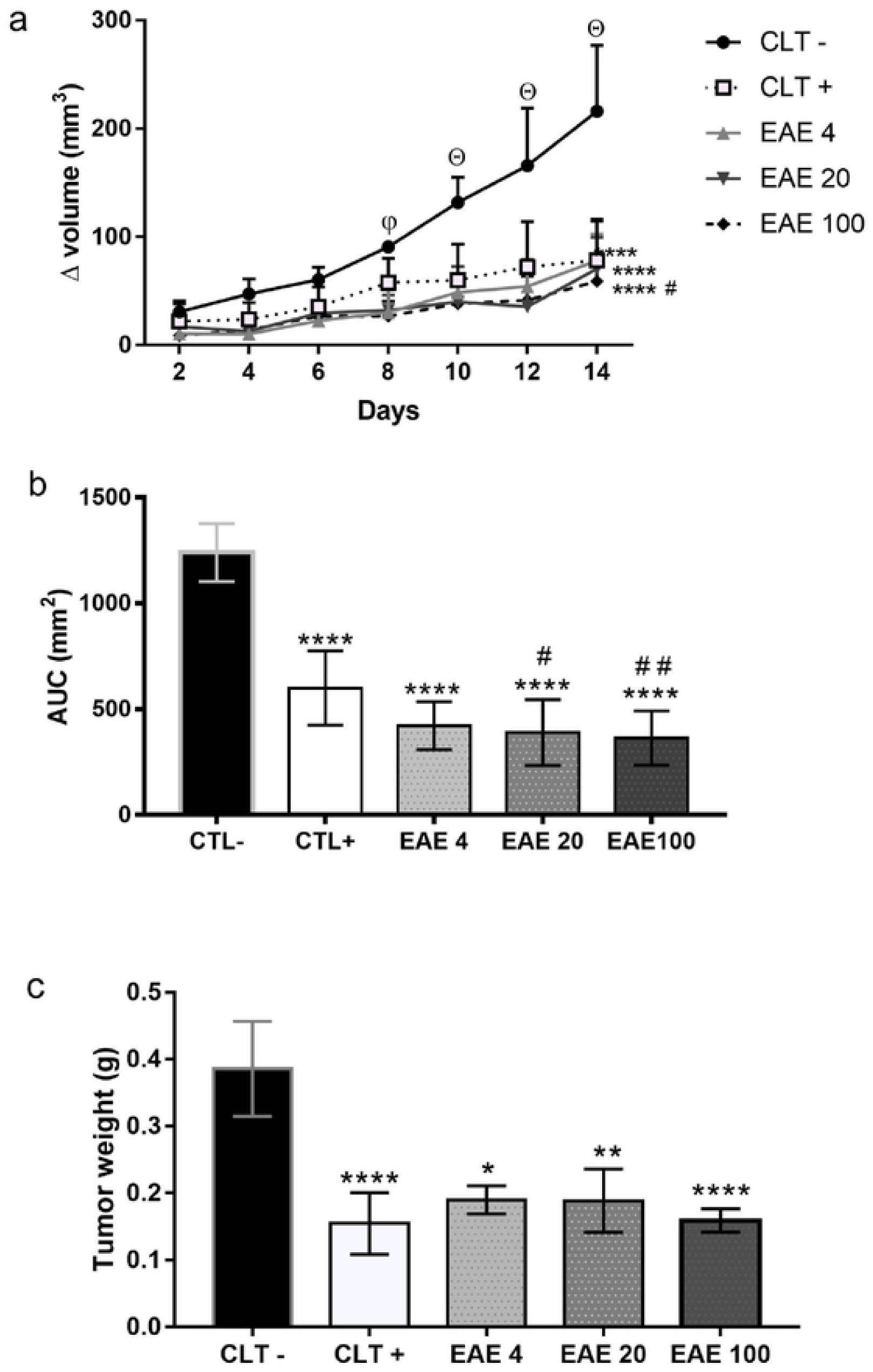
Effect of *Penicillium purpurogenum* extract on the development of Ehrlich’s solid tumor. (a) The kinetic of paw volume inoculated with Ehrlich’s tumor followed by intraperitoneal treatment with phosphate buffer solution (CTL-), cyclophosphamide 25mg/kg (CTL +) and extracts with a concentration of 4 mg/kg (EAE4), 20 mg/kg (EAE20) and 100 mg/kg (EAE10) at 24h intervals, with the animals being euthanized on day 15. (b) Area under the curve (AUC) calculated from the volume growth kinetics. (c) The average weight of the paws with a tumor was determined in the groups after treatment. Valeus expressed as mean ± standard error of means (S.E.M.) deviation and analyzed by the analysis of variance (one-way or two-way ANOVA) with *p<0.05, ** p <0.005, **** p <0.0001 in relation to the negative control (CTL-), #p < 0.05, ## p <0.01 when compared to positive control (CTL +) and Θ p<0.05 difference in tumor growth for the other groups and φ shows that, on the eighth day, only the extract inhibited tumor growth.

### Histological results

Histopathological analysis showed the presence of tumor masses in foot pad of animals from groups with Ehrlich’s solid tumor. The tumour masses exhibited high cellularity and different growth patterns, central, moderate to very high pleomorphism with bizarre nuclear forms, presence of one or more nucleoli and heterogeneous chromatin patterns. The cytoplasm was eosinophilic, abundant, with poorly defined limits. Occasional multinucleated giant cells (associated with pattern 3 pleomorphism) and up to 2 mitosis figures per 400x field were identified. Groups with tumor induction presented peritumoral, multifocal to coalescent lymphohistiocytic inflammatory infiltrate with variable intensity Areas of multifocal liquefaction necrosis, with accumulation of eosinophilic and amorphous cellular debris associated with neutophilic infiltrate were also observed. Interestingly, the extract showed an inflammatory process induced by the minor tumor in the groups treated with 20 and 100 mg/kg when compared to the negative control. The extract also demonstrated to have smaller areas of tumor necrosis, mitotic activity and invasion when compared to the negative control (Figure 3 and Table 1).

**Table 1.** Ehrlich tumor histopathological scores (mean ± SD) of groups treated with saline (CTL-), cyclophosphamide (CTL+) or *Penicillium purpurogenum* ethyl acetate extract at 4mg/Kg (EAE4), 20mg/Kg (EAE20) and 100 mg/Kg (EAE100).

|  | Pleomorphism | Necrosis | Mitosis<br>Figures | Inflammation | Invasion |
| --- | --- | --- | --- | --- | --- |
| **CLT-** | 2.4±0.511 | 2.4±0.843 | 1±1.155 | 2.8±0.421 | 2±0.666 |
| **CLT+** | 0.2*±0.421 | 0.2*±0.421 | 0*±0.000 | 1*±1.333 | 0.6*±0.843 |
| **EAE.4** | 2.2±1.122 | 1.2±1.033 | 1±1.155 | 1.4±1.075 | 1±0.667 |
| **EAE.20** | 2±1.155 | 0.6*±1.265 | 0.2±0.421 | 0.8*±0.788 | 1±0.667 |
| **EAE.100** | 0.8*±1.033 | 0.2*±0.421 | 0*±0.000 | 0.6*±0.843 | 0.4*±0.843 |

Scores: 0 (absent), 1 (weak), 2 (moderate), 3 (intense); the result was calculated by the mean of the scores; \* p <0.05 in relation to the negative control (CLT-). N = 5/group.

**Fig. 3.**
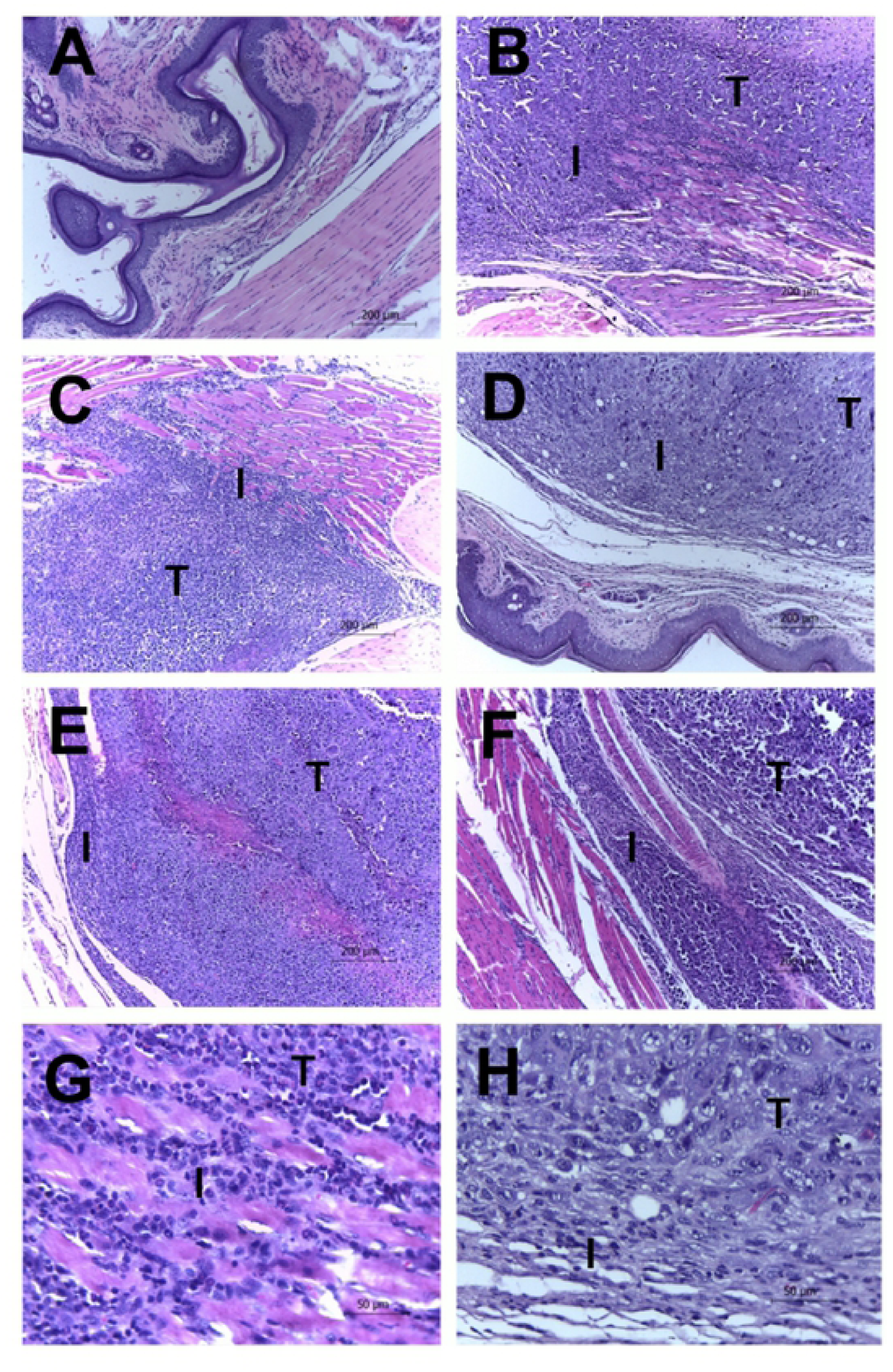
Tumor and leukocyte infiltration in Ehrlich tumors with and without treatments. Leukocyte infiltrates in Ehrlich tumors. Swiss mice were inoculated in the paw with 2×10^6^ Ehrlich tumor cells and were treated daily with EAE extract intraperitoneally. At the end of the fifteen days of treatment, the animals were euthanized and their feet amputated, weighed and fixed. Histological sections were stained with Hematoxylin-Eosin. In the photos it is possible to see the tumor cells (indicated by letter T) and the inflammatory infiltrate (indicated by letter I) present in the paws of the Sham **(A)**, CLT- **(B)**, CLT + **(c)** group (100x total magnification). The animals treated with the extract showed a decrease in the infiltrate and necrosis in the EAE doses EAE 4 **(D)**, EAE 20 **(E)** and EAE 100 **(F)** (100x total magnification). The Sham group is shown on panel **G** and animals treated with extract in the EAE 100 dose on panel **H** (400x total magnification).

### *P. purpurogenum* extract induced immunomodulatory effects

Treatment with EAE at doses of 20 mg/kg and 100 mg/kg presented low number of cells in popliteal lymph nodes (9.6×10^4^±1.0 cells/mL and 4.4×10^4^±0.34 cells/mL, respectively) when compared with negative control (326×10^4^±40.83 cells/mL). The similar results were observed in cyclophosphamide-treated animals (8.0×10^4^±1.88 cells/mL) (Figure 4a). In spleen cells number account, the negative control group (72.7×10^7^±7.06 cells/mL) showed high cellularity when compared to the animals of Sham group (1.6 ×10^7^±0.12 cells/mL), however this difference was not observed in animals inoculated with the tumor and treated with cyclophosphamide (0.7×10^7^±0.02 cells/mL) or with extract doses 20 mg/kg (1.6×10^7^±0.14 cells/mL) and 100 mg/kg (1.2×10^7^±0.04 cells/mL) (Figure 4b). The marrow bone cellularity showed that the cyclophosphamide (12.3×10^5^±3.13 cells/mL) and the extract treatment (61.0×10^5^±8.35 cells/mL, 17.6×10^5^±1.20 cells/mL and 4.0×10^5^±1.08cells/mL, respectively) prevented the high level of cellularity in associated with Ehrlich’s tumor, since the groups without tumor (sham: 4.6×10^5^±0.77 cells/mL and EAE20: 15.44×10^5^±5.04 cells/mL) had fewer cells than negative control group (189.0×10^5^±23.12 cells/mL) (Figure 4c). The results showed that extract’s effect was dose-dependent.

**Fig. 4.**
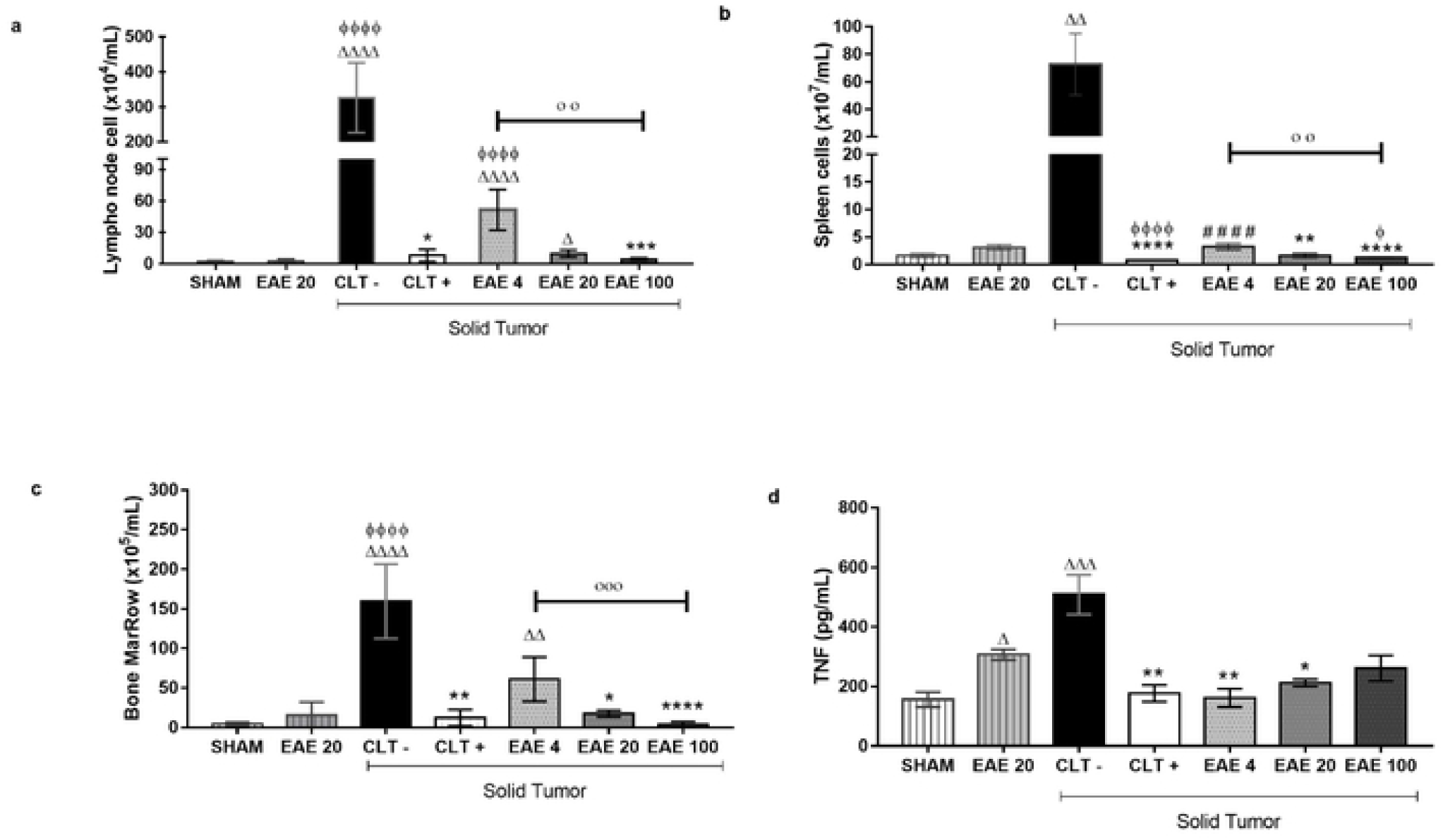
Immunomodulatory effects of *Penicilium purpurogenum* ethyl acetate extract. **(a, b e c)** The results were expressed as the mean ± standard deviation of the total lymph node cell count, bone marrow, splenocytes obtained from the group without tumor treated with saline solution (Sham), from the groups with solid Ehrlich tumor treated intraperitoneally with extracts at doses respectively 4, 20 and 100mg/kg (EAE4, EAE20 and EAE100), positive and negative control administered cyclophosphamide (CLT +) and e with saline solution (CTL-) respectively. **(d)** Blood serum was used for quantification of tumor necrosis factor (TNF-α). The data were submitted to statistical analyzes by method the Kruskal-Wallis and Dunn multiple comparison test, with significance of Δ p <0.05, ΔΔ p <0.005, ΔΔΔ p <0.0005, ΔΔΔΔ p <0.0001 in relation to Sham, ϕ p <0.05, ϕϕϕϕ p <0.0001 when compared to EA20 without tumor, #### p <0.0001 when compared to CLT+, * p <0.05, ** p <0.005, *** p <0.0005, **** p <0.0001 when compared to CLT-, ○○ p <0.0001, ○○○ p <0.0001 comparing the extracts with each other.

Cytokine quantification demonstrated that treatment was able to alter only the TNF levels (Supplementary Data 1). The animals with untreated tumours (CTL-) showed an even higher level in TNF-α concentrations (508.8±66.22 pg/mL) when compared with Sham group (156.0±25.14 pg/mL). The cyclophosphamide treatment significantly showed lower TNF-α levels (177.2±27.67 pg/mL) compared to the negative control group. Similarly, the extract at 4, 20 and 100mg/Kg doses also showed TNF-α low concentrations versus the group CLT- (161.7±30.22 pg/mL, 212.4±12.03 pg/mL and 261.4±43.16 pg/mL respectively) in animals with solid tumor. However, the extract treatment in animals without tumor showed high level of TNF-α seric concentration (306.9±17.93 pg/mL) when compared with normal animals (Figure 4d).

### Toxicity studies of *P. purpurogenum* EAE

Daily treatment with *P. purpurogenum* ethyl acetate extract in the solid Ehrlich tumor maintained the body weight of the tumor-inuculated animals (EAE4: 2.38±0.63 g; EAE20: 2.8±0.76 g and EAE100: 0.96±0.67 g) compared with CTL+ (−1.16±0.46 g) and CTL- (−0.24±0.73 g) (Figure 5a). The data showed that only the group receiving cyclophosphamide had a body weight reduction when compared with Sham group (1.58±0.96 g). There were no significant differences concerning serum AST and ALT levels in any of the groups analyzed, although CTL-animals showed increased average values compared with groups treated with the extract in the presence of the tumor (Table 2). Hepatic and renal hiltological difference was not found between the groups. The survival rate curve showed that saline (79.12%) and cyclophosphamide (85.72%) groups were statistically different, as expected. Importantly, tumor-animals groups treated with the extract showed 100% survival rate in all doses, being significantly different when compared to controls (Figure 5b).

**Table 2.** Effect of *Penicilium purpurogenum* ethyl acetate extract on ALT and AST serum levels (mean ± SEM) in all experimental groups.

|  | ALT (U/L) | AST (U/L) |
| --- | --- | --- |
| **Sham** | 125.2±103.5 | 125.2±103.5 |
| **EAE20\*** | 122.5±101.5 | 295.3±236.1 |
| **CLT-** | 207.9±181.8 | 438,3±328.5 |
| **CLT+** | 438.3±328.5 | 293±131.8 |
| **EAE4+Tumor** | 50.12±33.38 | 178.6±95.15 |
| **EAE20+Tumor** | 68.3±57.5 | 185.1±111.2 |
| **EAE100+Tumor** | 48.88±21.68 | 179.1±61.28 |

N=5 for each group. One-way ANOVA showed no statistical differences between groups. \*Tumor free group treated with extract in dose of 20mg/kg. The groups treated with saline (CTL-), cyclophosphamide (CTL+) and *Penicillium purpurogenum* ethyl acetate extract at 4mg/Kg (EAE4+Tumor), 20mg/Kg (EAE20+Tumor) and 100 mg/Kg (EAE100+Tumor).

**Fig. 5.**
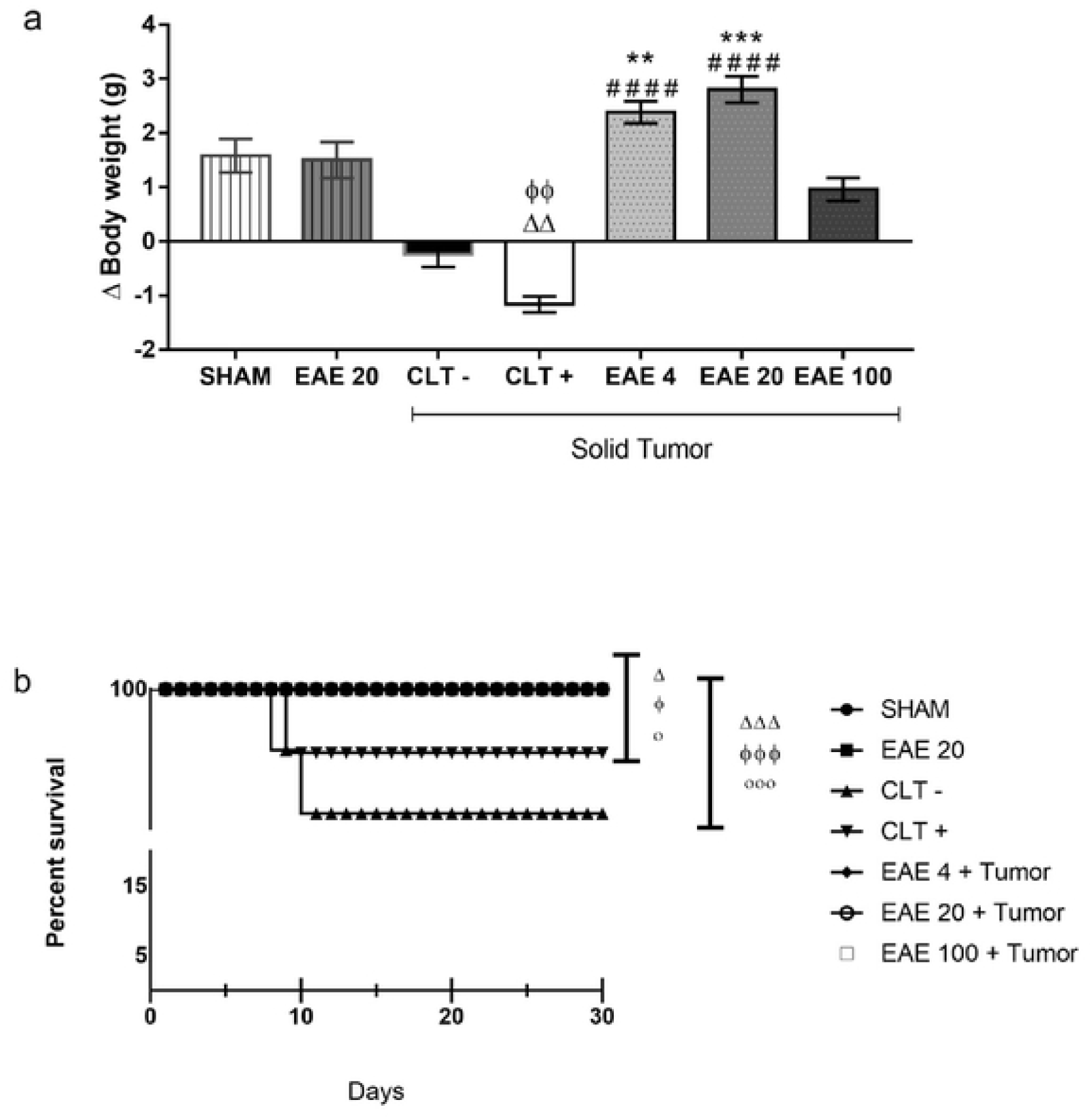
Effect of the toxicity of the extracellular extract of *Penicillium purpurogenum* ethyl acetate extract (EAE). The groups without tumor induction were treated with saline (sham), while groups with induction received extract at doses of 4, 20 and 100 mg/kg (EAE4, EAE20 and EAE100), saline (CTL-) and cyclophosphamide (CTL +), where we can see in the graphs: a) The animals’ body weight in the final treatment. (b) After the treatment of the animals, it remained under observation for thirty days. The data represent mean ± SEM. The difference was statistically analyzed by Kruskal-Wallis and Dunn’s multiple comparison test, with significance of Δ p <0.05, ΔΔp <0.005, ΔΔΔp <0.0005 in relation to Sham, ϕ p <0.05 ϕϕp <0.005, ϕϕϕp <0.0005 when compared to EA20 without tumor, ○ p <0.05 when compared to CLT + and ○○○ p <0.0005 when compared to CLT-.

## DISCUSSION

*P. purpurogenum* isolated from the marine environment is capable of secreting secondary bioactive metabolites with anticancer properties. In the present study, the extracellular extract of the fungus revealed the presence of macrophorin A, purpurogemutanthidine and purpurogemutanthin, corroborating previous findings by Tang et al. [32], Fang et al [29]. and He et al [33]. These compounds showed interesting anti-cancer activity in vitro against cervical cancer, gastric adenocarcinoma and breast cancer cells, while berkeleyacetal C [34,35,36] and rubratoxin B [37,38,39] showed anti-inflammatory activity and anticancer in vitro.

Meroterpenoids, which are formed by the mixed terpenoid-polyketide biosynthetic pathway, are an important class of compounds in the context of the development of new anticancer agents, due to their vast structural diversity and broad spectrum of bioactivities [40]. They are known to have activities against cancer through various mechanisms, such as the blocking of cell survival pathways, activity against oxidative stress, and induction of apoptosis [41]. When checking the presence of compounds with anticancer activity in vitro, we questioned whether the extract isolated from the marine environment and not described in the literature with activity in vivo against Erhlich’s tumor could be active.

The present study is the first to evaluate the *in vivo* anticancer effects of *P. purpurogenum*, employing mice inoculated with Ehrlich’s tumor cells. The results showed that extract compounds were efficient to reduce the development and weight of Ehrlich solid tumor.

Ehrlich’s tumor exhibits strong inflammatory phenomena, including edema and inflammatory infiltrates, which are believed to play an essential role in tumor growth in various types of cancer [42, 43, 44, 45, 46]. But the anti-inflammatory effects were reported by meroterpenoids [47] and were observed in this research showing that the extract reduces inflammation, reducing the tumor by an indirect route.

The extract inhibited tumor growth at the same level as the untreated group, since the behavior of the animals and the survival of the group treated with the extract was similar to the sham group, probably acting in tumor cell proliferation, migration, and invasion.

Two possibilities can inhibit tumor growth: the direct effect on the tumor microenvironment caused by macrophorin A, purpurogutididine and purpurogemutanthin identified in the extracts that result in smaller tumor size. In addition to an indirect effect, reducing inflammation which makes the tumor smaller, which may be associated with the presence of Berkeleyacetal C in the extract.

Other mereterpenoides shows inhibition tumor activities. Wang et al [37] studied Guajadial, one of natural dialdehyde meroterpenoids, that was capable to suppress tumor growth in human xenograft mouse models, with probably proliferation inhibition by the block the Ras/MAPK pathway.

In research carried out by Wan et al. [48] they demonstrate that mereterpenoids have anti-inflammatory and antioxidant activity that can allow the reduction of tumor growth. Li et al. [31] found that Berkeleyacetal C causes significant inhibition of the expression of inducible nitric oxide synthase (iNOS) and the production of nitric oxide by macrophages.

Berkeleyacetal C inhibits expression and secretion of major pro-inflammatory factors and chemokines, including tumor necrosis factor-α (TNF-α) interleukin-6 (IL-6), interleukin-1β (IL-1β), macrophage inflammatory protein −1α (PImax) −1α), and the monocyte chemotactic protein-1 (MCP-1) [29, 35].

Imunosupression cells can have pro-tumoral effect. Indeed, higher infiltration by Tregs is observed in tumor tissues, and their depletion augments antitumor immune responses in animal models [44].

There were also systemic effects, with dose-dependent reduction in the cells of the popliteal lymph node, spleen and bone marrow, when compared in the negative control group. This is in line with a marked low levels of TNF-α, which plays an important role in the beginning and applies the activation of adhesion molecules and expression of inflammatory mediators during inflammatory responses [49]. These pro-inflammatory mediators can cause damage to cells and tissues and also activate macrophages in various diseases associated with inflammation [50].

Increased levels of pro-inflammatory cytokines was previously associated with the development of Ehrlich’s tumor [51]. Our data also agree with those of Calixto-Campos et al [52], Aldubayan et al [53] and Harun et al [54], who studied the protective role of fungal extracts against inflammatory events induced by LPS in vitro and found that all the extracts inhibited the TNF-α expression. Taken together, these results show that the extracellular extract of *P. purpurogenum* has a potent anticancer activity in vivo against Ehrlich’s breast adenocarcinoma and this effect can be mediated, at least in part, by imnomodulatory mechanisms.

Another important set of observations from the present study concerns the safety of the extract. Remarkably, *P. purpurogenum’*s extracellular extract was able to improve mouse survival at all doses, compared with the negative control and even with cyclophosphamide, providing 100% survival at the end of the study. The extract preserved body weight of the animals, preventing the wasting syndrome that is often associated with cancer and intensified by chemotherapy [55,56,57,58,59], as previously reviewed [60,61].

These data suggest that *P. purpurogenum* extract may have potential for clinical use in combination therapies, to prevent the loss of body weight in cancer patients. Considering that fungal extracts often display hepatic toxicity, we evaluated ALT and AST serum levels and hepatic histology. Results did not demonstrate acute hepatic toxicity; instead, the extract not showed hepatic histological lesions associated with Ehrlich’s tumour and presented low serum levels of hepatic transaminases. In line with these observations, no changes were observed in renal histology. These results support the hypothesis that the extract has a favourable toxicological profile, although these observations should be confirmed and complemented by additional studies.

Thus, the results show that the extract inhibited tumor growth when compared to the negative control, at the same level as a standard therapy (cyclophosphamide), even when applied at the lowest dose.

Interestingly, the activity of the extract occurred before the cyclophosphamide, suggesting an intense anti-tumour activity *in vivo*. In addition, these findings correlate with morphological changes at the histological level, showed that *P. purpurogenum* presented low mitotic activity and invasive behaviour of tumour cells. The extract’s antitumour activity was associated with marked immunomodulatory effects.

## Conclusions

Overall, the present results indicate that the *P. purpurogenum* extract extract has potent *in vivo* anticancer activity against Ehrlich’s solid tumour, suggesting the presence of immunomodulatory mechanisms. Chemical components present in the extract may serve as lead compunds for the development of new compounds with antitumor effect and immune response modulators. The extract showed being well-tolerated and improved animal survival compared with cyclophosphamide, suggesting a favourable toxicity profile and potential applications in combination therapies. Further preclinical studies are required to clarify the potential uses of *P. purpurogenum* and to understand the production of bioactive metabolites in fungi for biotechnological applications.

## List of abbreviations

ALT: alanine aminotransferase
AST: aspartate aminotransferase
AUC: Area under the curve
CTL: negative control
CTL +: positive control
EAE: ethyl acetate extract
IFN-γ: Interferon gamma
iNOS: inducible nitric oxide synthase
IL-1β: interleukin-1β
IL-2: interleukin-2
IL-4: interleukin-4
IL-6: interleukin-6
IL-10: interleukin-10
IL- 17a: interleukin-17a
MCP-1: monocyte chemotactic protein-1
PImax-1α: macrophage inflammatory protein −1α
SEM: standard error of means
TNF-α: alpha Tumor Necrosis Factor

## Ethics approval and consent to participate

Animal studies were approved by ethical and research committee of Universidade Federal do Maranhão. This article does not describe any studies with human participants performed by any of the authors.

## Consent for publication

All the authors have consented for publication

## Availability of data and materials

The datasets supporting the conclusions of this article are included within the article and its additional files.

## Authors’ contributions

AMT, MDSBN and APSAS designed the concept and experiments under study. AMT and GFBB preparation of the fungus extract. APSAS and FRFN Implementation and standardization of the Erlich model. AMT, LPPS, SJAG, ALAB, GXS and APSAS performed the experiments. RMGC and GXS slide reading in the histological test. CJMT, AMT and FAS carried out the chemical characterization of the extract. FAS performed the quantification cytokines by FACS. AMT and APSAS Data curation; Formal analysis; Statistical analysis; Treatment of results in vivo and preparation of figures; Writing - original draft; proofreading and editing. MACNS: Writing - original draft; proofreading and editing. MDSBN Acquisition of resources; Project management; Supervision.

## Acknowledgements

We would like to thank to Fundação Oswaldo Cruz for providing the “*in vitro*” experimental procedures.

We would like to thank to Fundação de Amparo à Pesquisa e ao Desenvolvimento Científico e Tecnológico do Estado do Maranhão for financial support.

